# Mean-field models for EEG/MEG: from oscillations to waves

**DOI:** 10.1101/2020.08.12.246256

**Authors:** Á. Byrne, James Ross, Rachel Nicks, Stephen Coombes

## Abstract

Neural mass models have been actively used since the 1970s to model the coarse-grained activity of large populations of neurons. They have proven especially fruitful for understanding brain rhythms. However, although motivated by neurobiological considerations they are phenomenological in nature, and cannot hope to recreate some of the rich repertoire of responses seen in real neuronal tissue. Here we consider a simple spiking neuron network model that has recently been shown to admit to an exact mean-field description for both synaptic and gap-junction interactions. The mean-field model takes a similar form to a standard neural mass model, with an additional dynamical equation to describe the evolution of population synchrony. As well as reviewing the origins of this *next generation* mass model we discuss its extension to describe an idealised spatially extended planar cortex. To emphasise the usefulness of this model for EEG/MEG modelling we show how it can be used to uncover the role of local gap-junction coupling in shaping large scale synaptic waves.

## 1 Introduction

The use of mathematics has many historical successes, especially in the fields of physics and engineering, where mathematical concepts have been put to good use to address challenges far beyond the context in which they were originally developed. Physicists in particular are well aware of the “The Unreasonable Effectiveness of Mathematics in the Natural Sciences” [63]. One recent break-through in the field of large-scale brain modelling has come about because of advances in obtaining exact mean-field reductions of certain classes of coupled oscillator networks via the so-called Ott–Antonsen (OA) ansatz [46]. This is especially important because the mathematical step from microscopic to macroscopic dynamics has proved elusive in all but a few special cases. Indeed, many of the current models used to describe coarse-grained neural activity, such as the Wilson-Cowan [65], Jansen-Rit [23], or Liley [34] model are phenomenological in nature. Making use of the OA reduction Luke and colleagues [36, 52] were able to obtain exact asymptotic dynamics for networks of pulse-coupled theta neurons [13]. Although the theta-neuron model is simplistic, it is able to capture some of the essential features of cortical firing pattern, such as low firing rates. As such, this mean-field reduction is a candidate for a new type of cortical neural mass model that makes a stronger connection to biological reality than the phenomenological models mentioned above. The theta neuron is formally equivalent to the quadratic integrate-and-fire (QIF) model [33], a mainstay of many studies in computational neuroscience, e.g. [11]. Interestingly an alternative to the OA approach has been developed by Montbrió *et al*. [39] that allows for an equivalent reduction of networks of pulse-coupled QIF neurons, and establishes an interesting duality between the two approaches. In the OA approach the complex Kuramoto order parameter is a fundamental macroscopic variable and the population firing rate is function of the degree of dynamically evolving within-population synchrony. Alternatively in the approach of Montbrió *et al*. average voltage and firing rate couple dynamically to describe emergent population behaviour. Given that both approaches describe the same overall system *exactly* (at least in the thermodynamic limit of an infinite number of neurons) there must be an equivalence between the two macroscopic descriptions. Montbrió *et al*. have further shown that this relationship takes the form of a conformal map between the two physical perspectives. This correspondence is very useful when dealing with different types of neuroimaging modality. For example, when looking at power spectrograms from electro- or magneto-encephalograms (EEG/MEG), it is useful to contemplate the Kuramoto order parameter since changes in coherence (synchrony) of spike trains are likely to manifest as changes in power. On the other hand the local field potential recorded by an extracellular electrode may more accurately reflect the average population voltage. A model with a perspective on both simply by a mathematical change of viewpoint is not only useful for describing experimental data, it may also help the brain imaging community develop new approaches that can exploit a non-intuitive link between seemingly disparate macroscopic variables. Importantly, for this to be relevant to the real world some further features of neurobiology need to be incorporated, as purely pulsatile coupling is not expected to capture all of the rich behaviour seen in brain oscillations and waves. In particular synaptic processing and gap-junction coupling at the level of localised populations of neurons, and axonal-delays at the larger tissue scale are all well known to make a major contribution to brain rhythms, both temporal and spatio-temporal. Fortunately, these biological extensions, that generalize the initial theta-neuron and QIF network models with pulsatile coupling, are natural and easily accommodated. Work in this area has already progressed, e.g. with theoretical work by Laing on how to treat gap-junctions [30] and by Coombes and Byrne [10] on the inclusion of realistic synaptic currents (governed by reversal potentials and dynamic conductance changes). Recent work in [9] has also considered the inclusion of finite action potential speeds. In this paper we consider a synthesis of modelling work to date on developing a new class of mean-field models fit for use in complementing neuroimaging studies, and present some new results emphasising the important role of local gap-junction coupling in shaping brain rhythms and waves.

Even without the inclusion of gap-junction a first major success of this so-called *next generation* neural mass and field modelling approach has been in explaining the phenomenon of beta-rebound. Here a sharp decrease in neural oscillatory power in the 15 Hz EEG/MEG beta band is observed during movement followed by an increase above baseline on movement cessation. Standard neural mass models cannot readily reproduce this phenomenon, as they cannot track changes of synchrony within a population. On the other hand the next-generation models treat population coherence as fundamental, and are able to track and describe changes in synchrony in a way consistent with movement-related beta decrease, followed by an increase above baseline upon movement termination (post-movement beta rebound) [7]. Moreover, these models are capable of explaining the abnormal beta-rebound seen in patients with schizophrenia [8]. Beta decrease and rebound are special cases of event related synchrony/de-synchrony (ERS/ERD), as measured by changes in power at given frequencies in EEG/MEG recordings [47], and as such this class of model clearly has wider applicability than standard neural mass models that cannot describe ERD/ERS because their level of coarse-graining does not allow one to interrogate the degree of within-population synchrony. By merging this new dynamical model of neural tissue with anatomical connectome data it has also been possible to gain a perspective on whole brain dynamics, and preliminary work in [9] has given insight into how patterns of resting state functional-connectivity can emerge and how they might be disrupted by transcranial magnetic stimulation. Despite the success of the next generation models that include synaptic processing it is well to recognise the importance of direct electrical communication between neurons that can arise via gap-junctions. Without the need for receptors to recognise chemical messengers gap junctions are much faster than chemical synapses at relaying signals. The communication delay for a chemical synapse is typically in the range 1 − 100 ms, while that for an electrical synapse may be only about 0.2 ms. Gap-junctions have long been thought to be involved in the synchronisation of neurons [2, 3] and are believed to contribute to both normal [21] and abnormal physiological brain rhythms, including epilepsy [58, 38].

In section 2 we introduce the mathematical description for the microscopic spiking cell dynamics as a network of QIF neurons with both synaptic and gap-junction coupling. We present the corresponding mean-field ordinary differential equation model with a focus on the bifurcation properties of the model under variation of key parameters, including the level of population excitability and the strength of gap-junction coupling. A simple cortical model built from two sub-populations, one excitatory and the other inhibitory, is shown to produce robust oscillations via a Hopf bifurcation. The derivation of the macroscopic equations of motion is deferred to a technical appendix. This new class of neural mass model is used as a building block in section 3 to construct a continuum model of cortical tissue in the form of an integro-differential neural field model. Here, long-range connections are mediated by action potentials giving rise to space-dependent axonal delays. For computational ease we reformulate the neural field as a brain-wave partial differential equation, and pose it on idealised one- and two-dimensional spatial domains. A Turing analysis is performed to determine the onset of instabilities that lead to novel patterned states, including bulk oscillations and periodic travelling waves. These theoretical predictions, again with details deferred to a technical appendix, are confirmed against direct numerical simulations. Moreover, beyond bifurcation we show that the tissue model can support rich rotating structures, as well as localised states with dynamic cores. Finally, in section 4 we discuss further applications and extensions of the work presented in this paper.

## 2 Neural mass model

Here we describe a new class of neural mass model that can be derived from a network of spiking neurons. The microscopic dynamics of choice is the QIF neuron model, which is able to replicate many of the properties of cortical cells, including a low firing rate. In contrast to the perhaps more well studied linear or leaky IF model it is also able to represent the shape of an action potential. This is important when considering electrical synapses, whereby neurons directly “feel” the shape of action potentials from other neurons to which they are connected. An electrical synapse is an electrically conductive link between two adjacent nerve cells that is formed at a fine gap between the pre- and post-synaptic cells known as a gap junction and permits a direct electrical connection between them. They are now known to be ubiquitous throughout the human brain, being found in the neocortex [15], hippocampus [14], the inferior olivary nucleus in the brain stem [53], the spinal cord [48], the thalamus [22] and have recently been shown to form axo-axonic connections between *excitatory* cells in the hippocampus (on mossy fibers) [17]. It is common to view the gap-junction as nothing more than a channel that conducts current according to a simple ohmic model. For two neurons with voltages *v*_*i*_ and *v*_*j*_ the current flowing into cell *i* from cell *j* is proportional to *v*_*j*_ − *v*_*i*_. This gives rise to a *state-dependent* interaction. In contrast, chemical synaptic currents are better modelled with *event-driven* interactions. If we denote the *m*th firing time of neuron *j* by 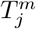 then the current received by neuron *i* would be proportional to 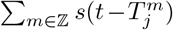 where *s* is a temporal shape that describes the typical rise and fall of a post synaptic response. This is often taken to be the Green’s function of a linear differential operator *Q*, so that *Qs* = *δ* where *δ* is a delta-Dirac spike. Through-out the rest of this paper we shall take *s*(*t*) = *α*^2^*t* exp(−*αt*)*H*(*t*), where *H* is a Heaviside step function. In this case the operator *Q* is second order in time and given by

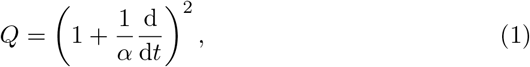

where *α^−^*^1^ is the time-to-peak of the synapse.

We are now in a position to consider a heterogeneous network of *N* quadratic integrate-and-fire neurons with voltage *v*_*i*_ and both gap-junction and synaptic coupling:

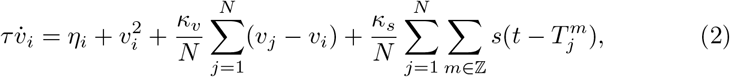

*i* = 1, …, *N*, with *v*_r_ ≤ *v*_*i*_ ≤ *v*_*th*_. Here, firing times are defined implicitly by 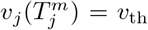. The network nodes are subject to reset: *v*_*i*_ → *v*_*r*_ at times 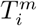. The parameter *τ* is the membrane time constant. The strengths of gap-junction and synaptic coupling are *κ*_*v*_ and *κ*_*s*_ respectively. The background inputs *η*_*i*_ are random variables drawn from a Lorentzian distribution with mean *η*_0_ and half width *γ*. The value of *η*_0_ can be thought of as setting the level of excitability, and *γ* as the degree of heterogeneity in the network. The larger *η*_0_ is, the easier it is to fire, and the larger *γ* is, the more dissimilar the inputs are. A schematic of a QIF network and its reduction to a neural field model is shown in Fig. 1, with details of the neural field formulation described next in section 3. The mean-field reduction of (2) can be achieved by using the approach of Montbrió *et al*. [39].

**Figure 1:**
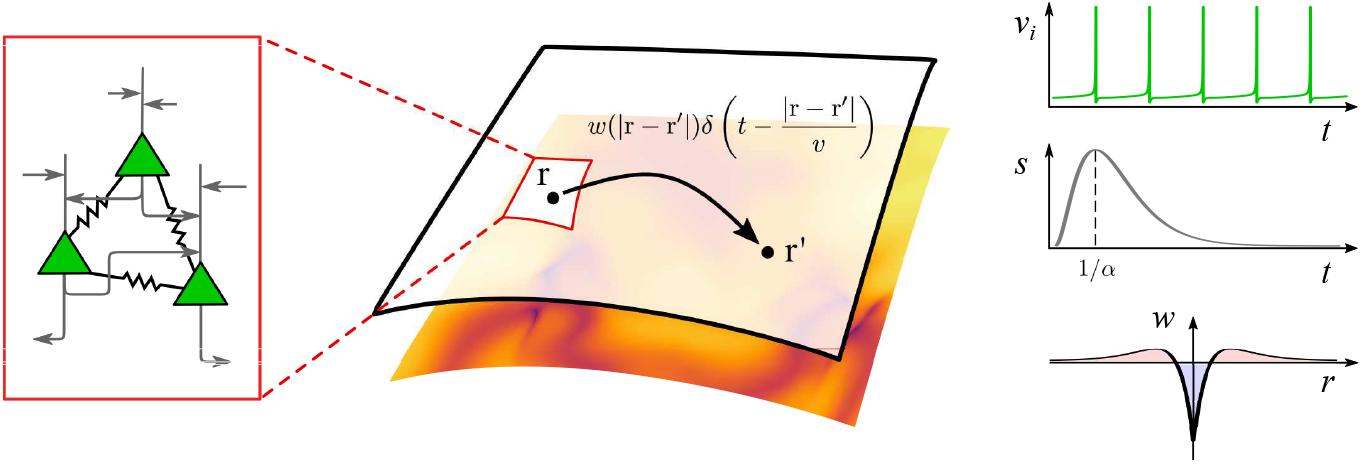
Model schematic. At each point in a two-dimensional spatial continuum there resides a density of QIF neurons whose mean-field dynamics are described by the triple (*R, V, U*), where *R* represents population firing rate, *V* the average membrane potential, and *U* the synaptic activity. The non-local interactions are described by a kernel *w*, taken to be a function of the distance between two points. The space-dependent delays arising from signal propagation along axonal fibres are determined in terms of the speed of the action potential *v*.

This is described in detail in Appendix A, and is valid for globally coupled cells in the thermodynamic limit *N → ∞*. The network behaviour can be summarised by the instantaneous mean firing rate *R*(*t*) (the fraction of neurons firing at time *t*), the average membrane potential 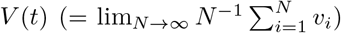, and the synaptic activity *U* (*t*). The synaptic activity *U* is driven by mean firing rate according to *QU* = *R*, with the mean-field dynamical equations for (*R, V*):

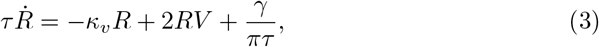

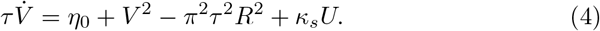

Interestingly this (*R, V*) perspective on population dynamics can be mapped to one that tracks the degree of within-population synchrony described by the complex Kuramoto order parameter *Z* according to the conformal map [39]:

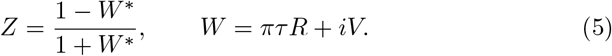

The corresponding dynamics for *Z* is given by equation (23) in Appendix A. Alternatively, one can evolve the model for (*R, V, U*) and then obtain results about synchrony |*Z*| by the use of (5).

### 2.1 Single population: bifurcation analysis

We first consider a single excitatory population (*κ*_*s*_ > 0). Unlike many traditional single population neural mass models, the activity of this next-generation model can oscillate in time (Fig. 2). Examining the profile of these oscillations, we observe that the peaks and troughs of the firing rate *R* and the synchrony |*Z*| roughly coincide. This indicates, rather unsurprisingly, that when a population is highly synchronised the population firing rate will be high.

**Figure 2:**
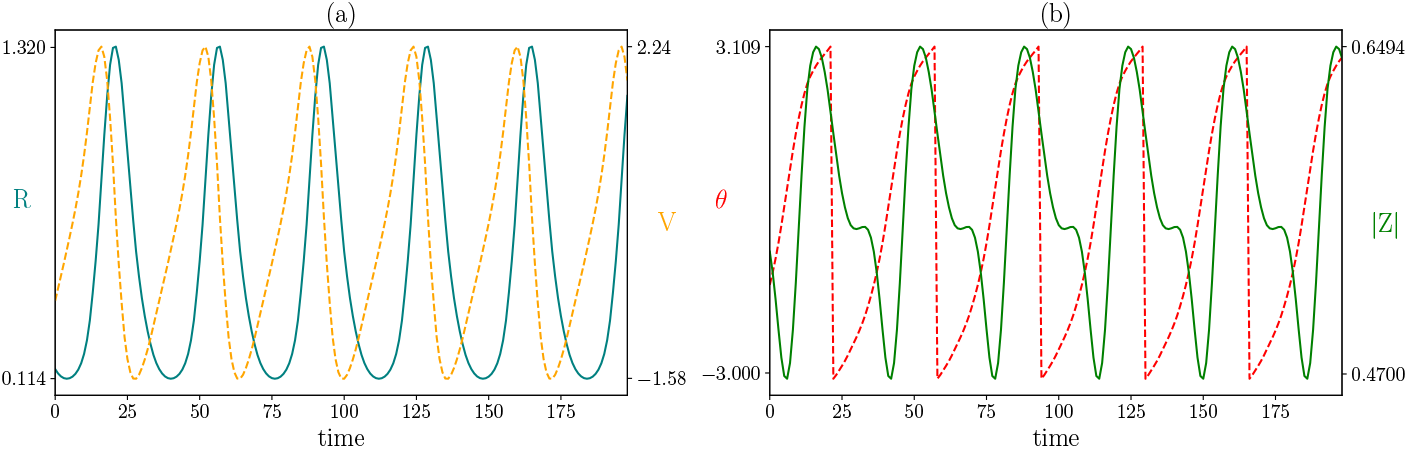
Single population dynamics: (a) Oscillations in the population firing rate *R* and average membrane voltage *V*, (b) Corresponding oscillations in the complex Kuramoto order parameter *Z* = |*Z*|e^*iθ*^, where |*Z*| reflects the degree of within-population synchrony, and *θ* a corresponding phase. Parameter values: *η*_0_ = 1, *κ*_*v*_ = 1.2, *κ*_*s*_ = 1, *τ* = 1, *α* = 1, *γ* = 0.5.

As the strength of gap-junction coupling *κ*_*v*_ is decreased the system undergoes a Hopf bifurcation and oscillations disappear (Fig. 3). Note that to the right of the Hopf bifurcation the amplitude of the oscillations increases with *κ*_*v*_. Increasing the level of excitability *η*_0_ also leads to oscillatory behaviour. A continuation of the Hopf bifurcation in *κ*_*v*_ and *η*_0_ is shown for different values of *γ* (Fig. 3). The system oscillates for parameter values to the right of these curves. Remembering that *γ* sets the level of heterogeneity, we note the window for oscillations gets smaller as the heterogeneity of network is increased.

**Figure 3:**
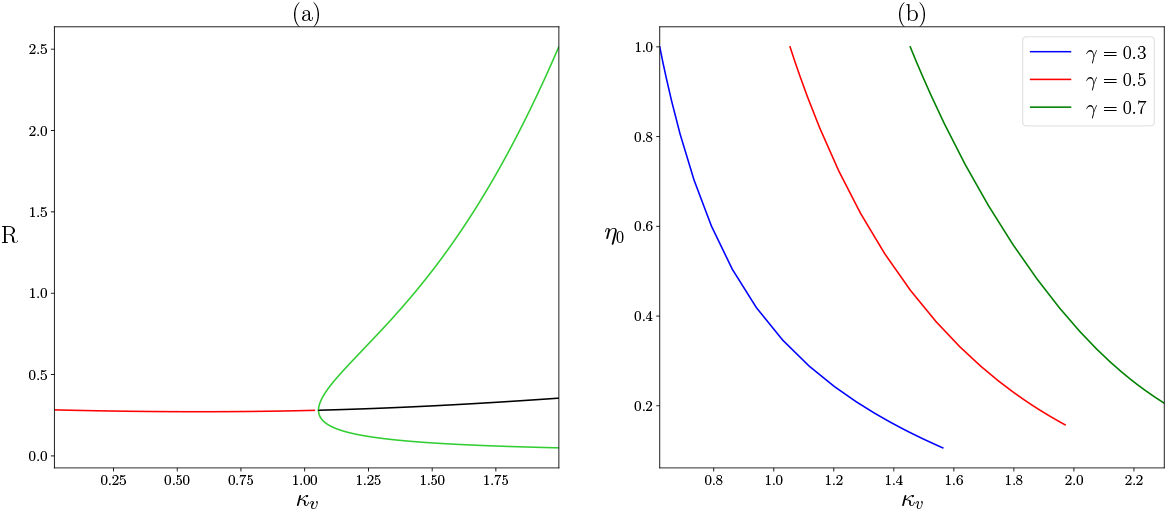
Single population bifurcation diagrams. (a) A Hopf bifurcation is found with an increase in the strength of gap junction coupling *κ*_*v*_, giving rise to limit cycle oscillations. Red (black) lines denote the stable (unstable) fixed point, while the green lines show the minimum and maximum of the oscillation. (b) A two parameter bifurcation diagram in the (*κ_v_, η*_0_)-plane tracing the locus of Hopf bifurcations for different values of *γ*. Oscillations emerge to the right of each curve. Parameter values: *η*_0_ = 1, *κ*_*s*_ = 1, *τ* = 1, *α* = 1, *γ* = 0.5.

### 2.2 Excitatory-inhibitory network: bifurcation analysis

The single population model can be easily extended to a two population network, consisting of an excitatory and an inhibitory population, labelled by *E* and *I* respectively. Synaptic coupling is present both within and between populations, while gap-junction coupling only exists between neurons in the same population. The augmented system of equations, describing the mean firing rate *R* and the average membrane potential *V* of each population, as well as 4 distinct synaptic variables *U* for each of the synaptic connections, is presented in Appendix B.

The excitatory-inhibitory network possesses a rich repertoire of dynamics. For example, it is possible to generate bursts of high frequency and high amplitude activity at a slow burst rate (Fig. 4). This pattern of activity is typical in epileptic seizures, e.g. [25]. Decreasing the gap-junction coupling strengths 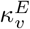 and 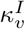 results in smoother lower amplitude oscillations, more in line with healthy brain oscillations. We note that 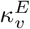 and 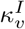 are not the only parameters that can change the profile of the oscillations; reducing 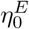 (the mean back-ground drive to the excitatory population) can also eradicate the seizure-like oscillations.

**Figure 4:**
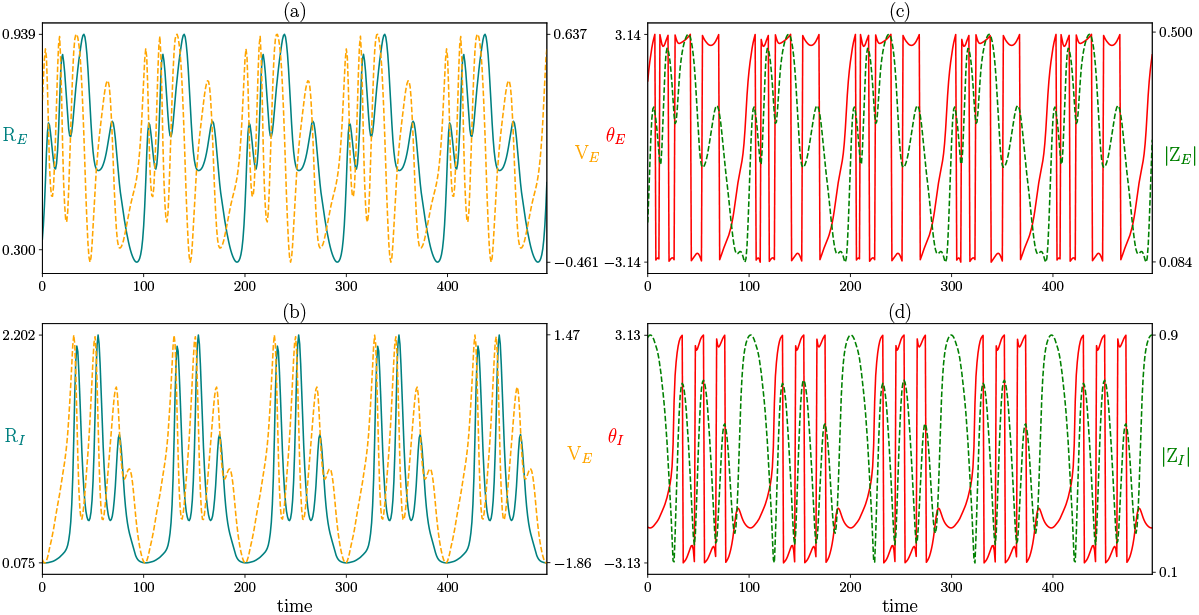
Excitatory-inhibitory network dynamics: (a) Oscillations in the excitatory population firing rate *R*_*E*_, as well as the average membrane potential *V*_*E*_. (b) Corresponding oscillations for the inhibitory population, *R*_*I*_ and *V*_*I*_. (c) Kuramoto order parameters for the excitatory population 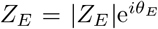. (d) Corresponding traces for *|Z_I_ |* and *θ*_*I*_ of the inhibitory population. Parameter values: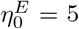, 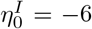, 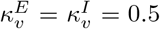, 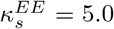, 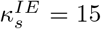, 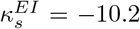,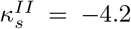, *τ*_*E*_ = 2*τ*_*I*_ = 1, *α*_*EE*_ = 1, *α*_*IE*_ = 1.4, *α*_*EI*_ = 0.7, *α*_*II*_ = 0.4, *γ*_*E*_ = *γ*_*I*_ = 0.5.

Next we examine the bifurcation structure of the excitatory-inhibitory network for different combinations of gap-junction coupling strengths 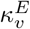 and 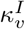 (Fig. 5). With no gap junction coupling in either population ((a) 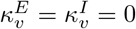, intermediate values of the mean background drive to the inhibitory population 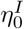 result in oscillatory behaviour. Switching on the gap junction coupling in the inhibitory population only ((b) 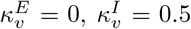, we observe oscillations for large 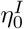 also. The firing rate of the excitatory population *R*_*E*_ performs low amplitude oscillations, while the firing rate of the inhibitory population *R*_*I*_ oscillates at a larger amplitude. Turning off the gap junction coupling in the inhibitory population but switching it on for the excitatory population ((c) 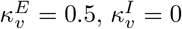, a different set of oscillatory solutions emerge for low 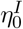. The amplitude of these oscillations is high for the excitatory firing rate, but low for the inhibitory population. Interestingly, the two oscillatory solutions co-exist for *η*_0_ 10 to 5. Jansen and Rit [23] demonstrated that transitions between seizures and healthy brain activity could be viewed as transitions between co-existing oscillatory solutions. A similar approach for this model (without gap junctions) can be found in [9]. With gap junctions switched on in both populations ((d) 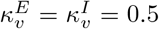, the 3 oscillatory solution branches exist and we see oscillations for nearly all values of 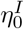.

**Figure 5:**
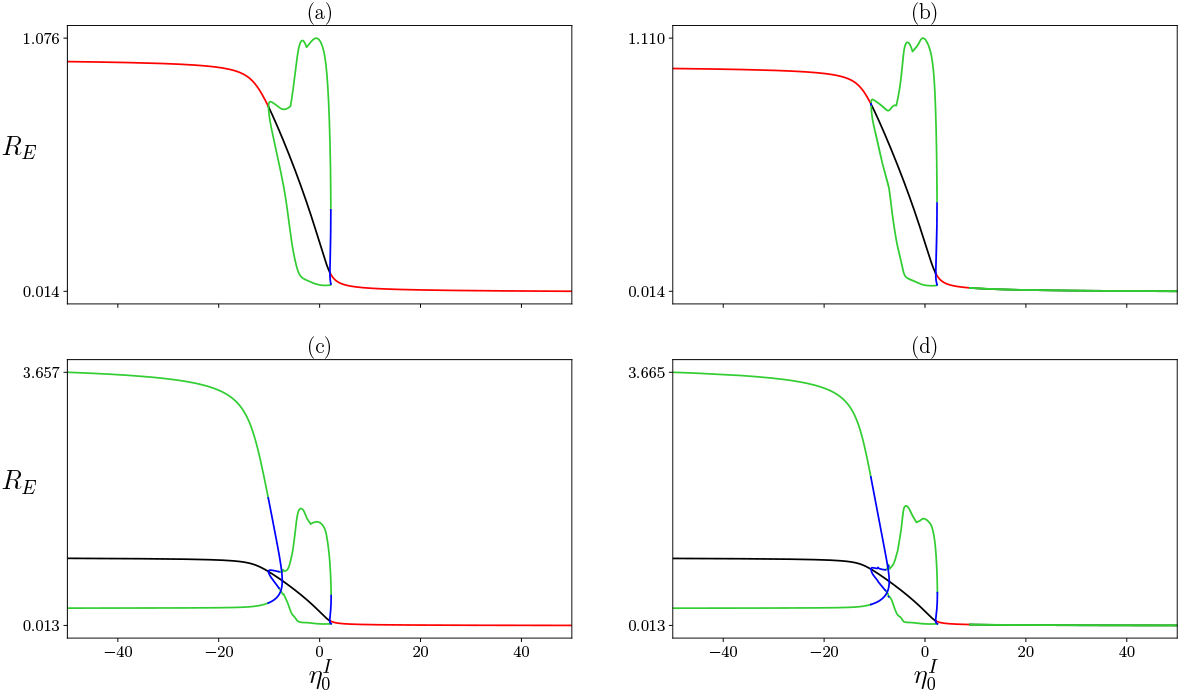
Two population bifurcation diagrams: Continuations in the mean background drive to the inhibitory population 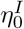 for different combinations of gap junction coupling strengths 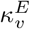 and 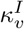. (a) No gap junction coupling, 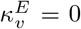, 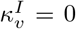 (b) Gap junctions in the inhibitory population only, 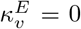, 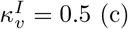 (c) Gap junctions in the excitatory population only, 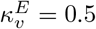, 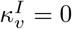 (d) Gap junction coupling in both populations, 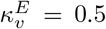, 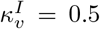. Other parameter values: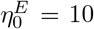, 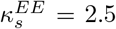, 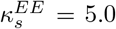, 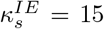, 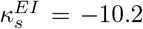, 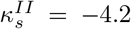, *τ*_*E*_ = 2*τ*_*I*_ = 1, *α*_*EE*_ = 1, *α*_*IE*_ = 1.4, *α*_*EI*_ = 0.7, *α*_*II*_ = 0.4, *γ*_*E*_ = *γ*_*I*_ = 0.5, *θ*_*E*_ and *θ*_*I*_.

With a good understanding of how the spatially clamped system behaves, we move on to the spatially extended neural field model.

## 3 Neural field model

Brain waves are inherently dynamical phenomena and come in a vast variety of forms that can be observed with a wide range of neuroimaging modalities. For example, at the mesoscopic scale it is possible to observe a rich repertoire of wave patterns, as seen in voltage-sensitive dye imaging data from the primary visual cortex of the awake monkey [41], and local field potential signals across the primary motor cortex of monkeys [50]. At the whole brain scale they can manifest as EEG alpha oscillations propagating over the scalp [20], and as rotating waves seen during human sleep spindles with intracranial electrocorticogram recordings [40]. Waves are known to subserve important functions, including visual processing [51], saccades [66], and movement preparation [50]. They can also be associated with dysfunction and in particular epileptic seizures [38]. Computational modelling is a very natural way to investigate the mechanisms for their occurrence in brain tissue, as well as how they may evolve and disperse [18, 19, 35].

The study of cortical waves (at the scale of the whole brain) is best advanced using a continuum description of neural tissue. The most common of these are referred to as neural fields, and are natural extensions of neural mass models to incorporate anatomical connectivity and the associated delays that arise through wiring up distant regions using axonal fibres. The study of waves, their initiation, and their interactions is especially pertinent to the study of epileptic brain seizures and it is known that gap junctions are especially important in this context [38]. Phenomenological neural field models with gap-junction coupling have previously been developed and analysed by Steyn-Ross *et al*. [56, 54], and more principled ones derived from *θ*-neuron models by Laing [29, 30]. In the latter approach it was necessary to overcome a technical difficulty by regularising the shape of the action potential. However, with the approach used in section 2 this is not necessary and the neural field version of (3)–(4) is constructed by replacing full temporal derivatives by partial temporal derivatives and replacing the temporal dynamics for *U* with the dynamics *QU* = ψ, where ψ denotes a spatio-temporal drive. For example, in the plane we might consider

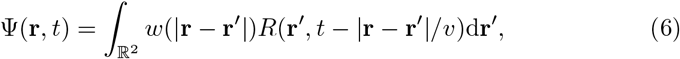

where **r** ∈ ℝ^2^ and *v* represents the speed of an action potential, as illustrated in Fig. 1. Typical values for cortico-cortical axonal speeds in humans are distributed, and appear to peak in the 5 − 10 m/s range [44]. Here, *w* represents structural connectivity as determined by anatomy. For example, long-range corticocortical interactions are predominantly excitatory whilst inhibitory interactions tend to be much more short-ranged, suggesting a natural choice for the shape of *w* as an inverted Mexican hat. A similar equation would hold in one spatial dimension. In this section we shall work with the explicit choice *w*(*x*) = (|*x*| − 1)e^−|*x*|^ in 1D and *w*(*r*) = (*r*/2 − 1)e^−*r*^/(2*π*) in 2D. For convenience we have chosen spatial units so that the scale of exponential delay is unity, though note that typical values for the decay of excitatory connections between cortical areas (at least in macaque monkeys) is ~ 10mm [37]. Both of the above kernel shapes have an inverted wizard hat shape and are balanced in the sense that the integral over the whole domain is zero. They also allow for a reformulation of the neural field model as a partial differential equation, as detailed in Appendix C. The resulting brain-wave equation is very amenable to numerical simulation using standard (e.g. finite difference) techniques. Before we do this, it is first informative to determine some of the patterning properties of the neural field model using a Turing instability analysis. Below we outline the results of the analysis and discuss the ensuing patterns for the neural field model in both 1D and 2D.

### 3.1 One spatial dimension

Turing instability analysis, originally proposed by Alan Turing in 1952 [57], is a mechanism for exploring the emergence of patterns in spatio-temporal system, including neural fields. Similar to the bifurcation analysis for the neural mass model, it allows us to determine the parameter values for which oscillations and patterns occur. Bulk oscillations, whereby synchronous activity across the spatial domain varies uniformly at the same rate, emerge at a Hopf bifurcation. Static patterns, which do not change with time, emerge at a Turing bifurcation. Dynamic patterns, that oscillate in time and space, emerge at a Turing-Hopf bifurcation.

The 1D neural field model, given in Appendix B by (36), supports both bulk oscillations and spatio-temporal patterns. Using the inverted wizard hat connectivity kernel (long-range excitation and short-range inhibition), we find Hopf and Turing-Hopf bifurcations (Fig. 6 left). See Appendix D for details of the analysis. For the chosen parameter values and weak gap-junction coupling (*κ*_*v*_ ≲ 0.8), the spatially-uniform steady state is always stable and neither patterns or oscillations exist. Increasing the mean background drive *η*_0_ moves the Hopf and Turing-Hopf curves down in the *v*-*κ*_*V*_ plane, allowing for oscillations and patterns in the absence of gap junctions (*κ*_*v*_ = 0). For slow action potential speeds (*v* ≲ 1), the system first undergoes a Hopf bifurcation as *κ*_*v*_ is increased and bulk oscillations emerge (Fig. 6 I). For faster action potential speeds (*v* ≳ 1), the Turing-Hopf bifurcation occurs before the Hopf, and we see periodic travelling and standing waves between the two bifurcations (Fig. 6 II). For lower action potential propagation speeds *v* we see standing waves close to the Turing-Hopf bifurcation (Fig. 6 III).

**Figure 6:**
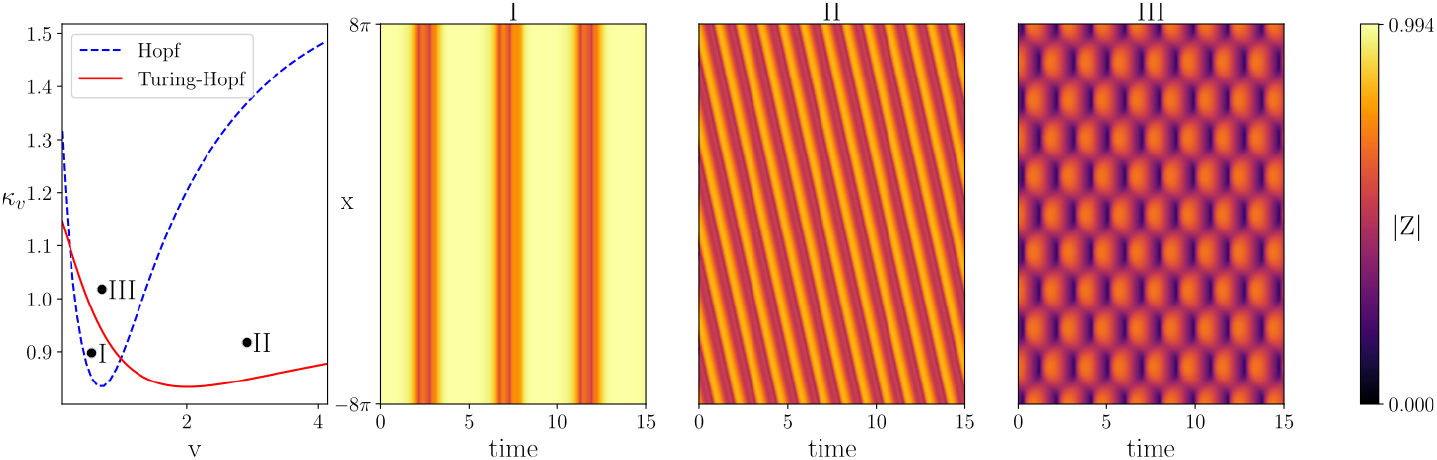
Turing instability analysis for the one-dimensional neural field model. The left panel shows the Hopf and Turing-Hopf curves as a function of the action potential speed *v* and gap junction coupling strength *κ*_*v*_. Above these curves patterned states emerge. The three right hand panels show simulations near Hopf, and two Turing-Hopf points: (I) Bulk oscillation with *v* = 0.7, *κ*_*v*_ = 0.9, (II) Periodic travelling wave with *v* = 3, *κ*_*v*_ = 1.2, (III) Standing wave with *v* = 0.9, *κ*_*v*_ = 1. Other parameter values: *η*_0_ = 0.3, *κ*_*s*_ = 5, *τ* = 1, *α* = 3, *γ* = 0.5.

To assess the role of gap junctions, we fixed the action potential speed *v* = 1 and explore the dynamics of the synchrony variable *|Z|* for different gap-junction coupling strengths *κ*_*v*_ (Fig. 7). For weak gap-junction coupling (I), there is a regular standing wave and the level of synchronisation oscillates between roughly 0.1 and 0.68. As *κ*_*v*_ is increased the amplitude of the oscillations increases, with the peak in synchrony reaching to about 0.85 for *κ*_*v*_ = 1.4 (II), 0.96 for *κ*_*v*_ = 1.7 (III) and 0.98 for *κ*_*v*_ = 2 (IV). Increasing *κ*_*v*_ further deforms the periodic standing wave pattern (III) and for *κ*_*v*_ = 2 the spatial pattern breaks down entirely (IV) in favour of synchronised spatially uniform oscillations. Note also that for strong gap-junction coupling (IV) the fluctuations in synchrony no longer reach down to ǀ*Z*ǀ ~ 0, suggestive of highly synchronised seizure like activity.

**Figure 7:**
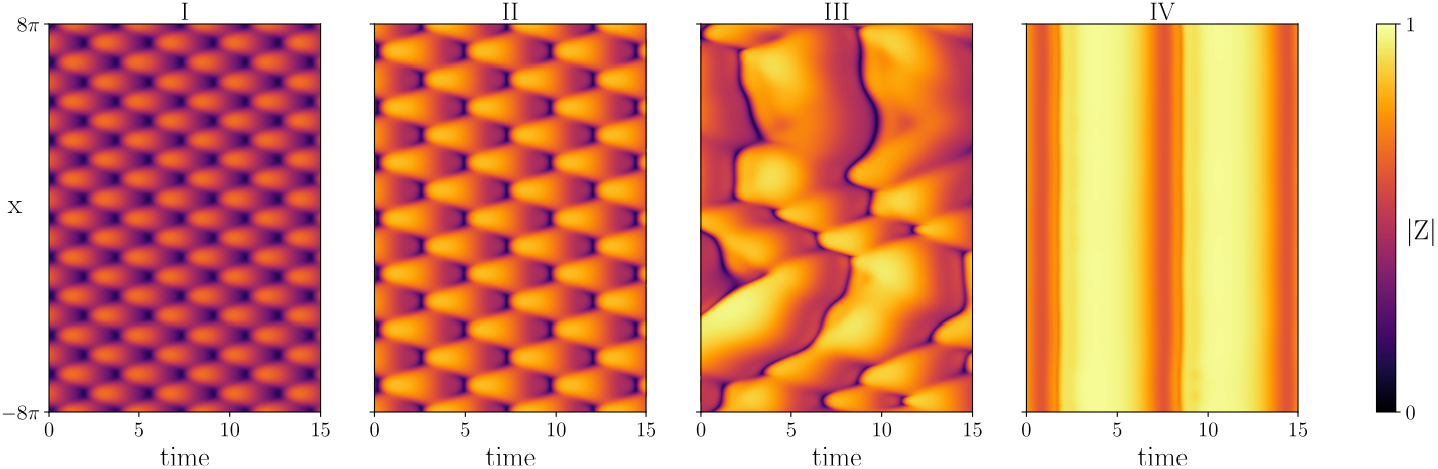
Simulations of the one-dimensional neural field model under variation in *κ*_*v*_: (I) Standing wave with *κ*_*v*_ = 1, (II) Standing wave (increased synchrony) with *κ*_*v*_ = 1.2, (III) Irregular dynamics (deformation of standing wave) with *κ*_*v*_ = 1.7, and (IV) Spatially synchronous oscillation with *κ*_*v*_ = 2. Other parameter values: *v* = 1, *η*_0_ = 0.3, *κ*_*s*_ = 5, *τ* = 1, *α* = 3, *γ* = 0.5.

For a standard wizard hat coupling kernel (long-range inhibition and short-range excitation) the neural field model can undergo a Turing bifurcation, as well as Hopf and Turing-Hopf bifurcations (see Supplemental information 1 panel (a)). Changing the sign of the synaptic coupling strength *κ*_*S*_ changes the coupling to long-range inhibition and short-range excitation. When Turing and Hopf instabilities occur simultaneously, interesting patterns emerge. In particular, we see stationary bumps where the activity at the centre of the bump oscillates in both space and time (see Supplemental information 1 panel (b)). This is akin to a chimera, some parts of the tissue are synchronised with one another while the bumps are incoherent. We will discuss the two dimensional version of such patterns in more detail below.

### 3.2 Two spatial dimensions

A Turing analysis was also performed for the 2D neural field equation, given in Appendix B by (35), and a very similar bifurcation structure was found when the mean background drive *η*_0_ = 0.5 (see Supplemental information 2 panel (a)). Increased *η*_0_, the Hopf and Turing-Hopf curves move down in the *v*-*κ*_*v*_ plane, and they switch, such that the Turing-Hopf occurs first for low action potential propagation speeds *v* as the gap junction coupling strength *κ*_*v*_ is increased (see Supplemental information 2 panel (b)). As expected, close to the Hopf bifurcation the activity of the tissue oscillates in time, but no spatial pattern emerges (see Supplemental information 3). Near the Turing-Hopf bifurcation we see both travelling and standing waves, depending on initial conditions (Supplemental information 4 (planar waves), 5 (radial waves) and 6 (standing waves)). Away from bifurcation, more interesting patterns emerge.

We fix the action potential speed *v* = 2 and vary the gap junction coupling strength *κ*_*v*_ to assess how gap junction coupling affects patterning. For weak gap junction coupling, we observe rotating waves with source and sink dynamics where the waves collide with each other (Fig. 8). The domain shown contains 8 rotating cores. Periodic boundary conditions were used. Hence, the cores at the edge of the domain wrap around to those on the other side. Supplemental information 7 shows the temporal evolution of the synchrony variable *Z*, from which the cores and rotations are readily observed. The direction of rotation alternates, such that every second core rotates clockwise/anti-clockwise.

**Figure 8:**
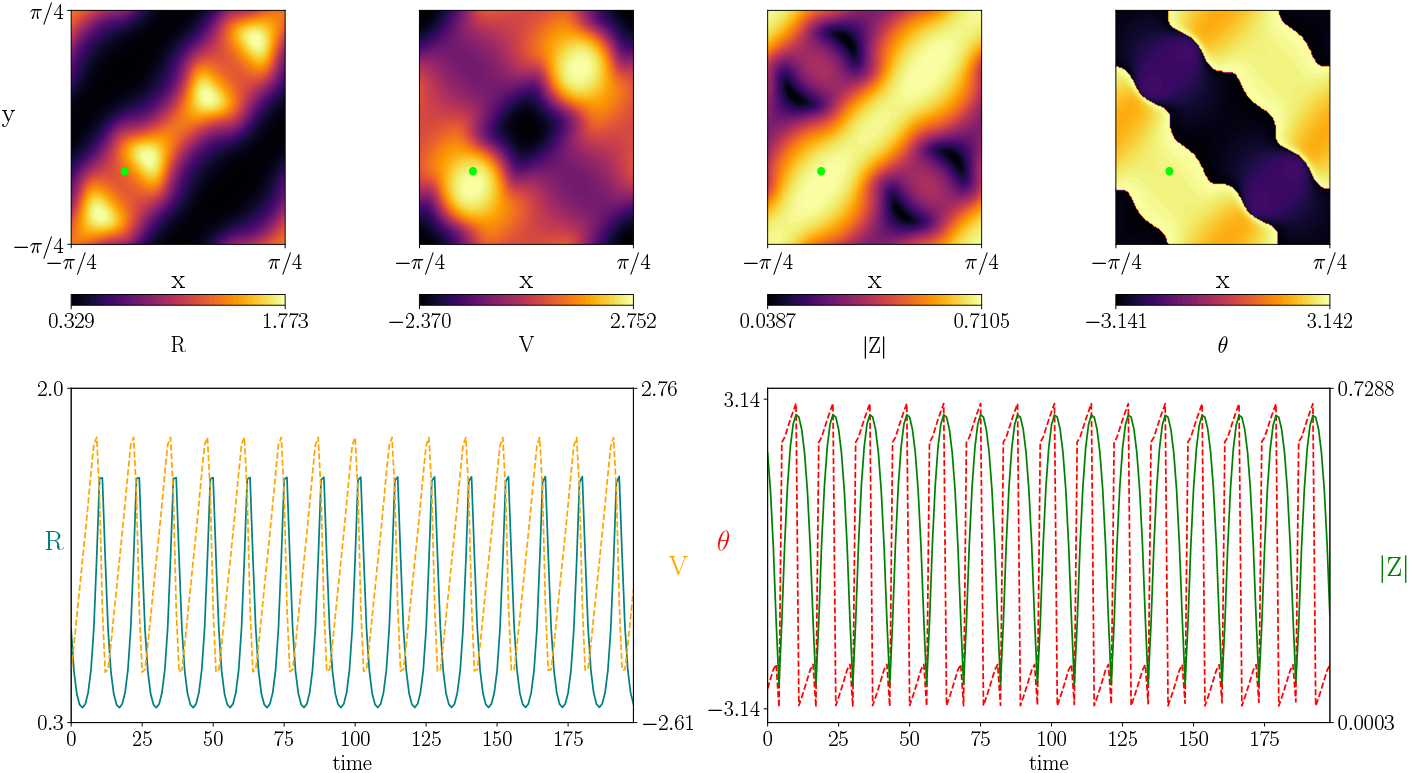
Simulations of the two-dimensional neural field model showing that, beyond a dynamic Turing instability, rotating waves with source and sink dynamics may emerge. Top: a snapshot of a patterned state in the (*R, V*) and (ǀ*Z*ǀ, *θ*) variables. Bottom: the corresponding time-series for the point marked by the small green circle in the top panel. A movie illustrating how this pattern evolves in time is given in Supplemental information 7. Parameter values: *v* = 2, *η*_0_ = 6, *κ*_*v*_ = 0.35, *κ*_*s*_ = 12, *τ* = 1, *α* = 5, *γ* = 0.5.

As the gap junction coupling strength *κ*_*v*_ is increased robust spirals emerge at the centre of the rotating cores. The spiral is tightly wound with a diffused tail of high amplitude activity that propagates into the rest of the domain and interacts with the other rotating waves (Fig. 9). The time course of a point close to the centre of a rotating core (green dot) depicts higher amplitude oscillations for the firing rate *R*, mean membrane potential *V* and level of synchronization *|Z|* when compared to the simulations for lower gap junction coupling strength *κ*_*v*_ (Fig. 8). In addition, the peaks in *R* are sharper and the minimum level of synchrony ǀ*Z*ǀ is substantially higher. The temporal evolution for the full tissue can be seen in Supplemental information 8.

**Figure 9:**
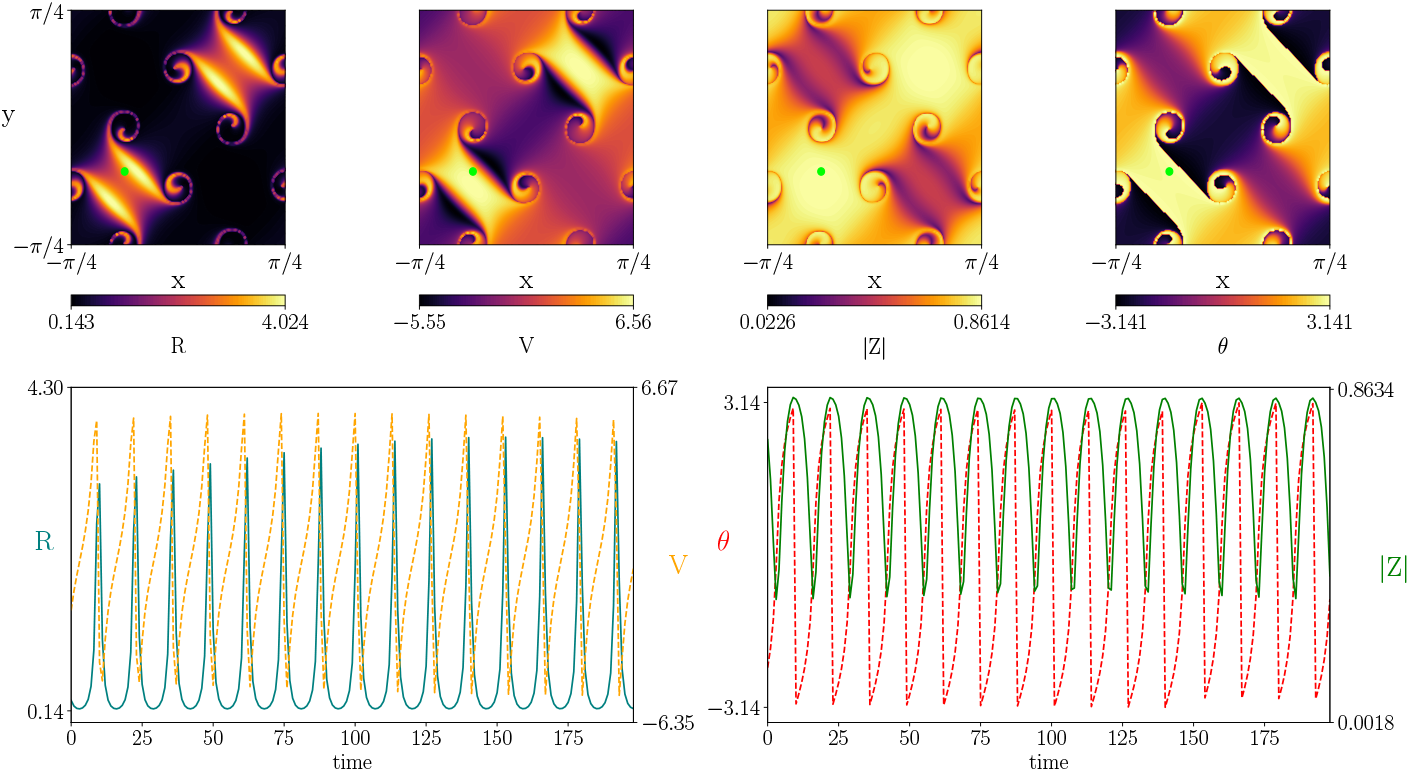
Simulations of the two-dimensional neural field model with moderate gap junction coupling strength. In this case robust spiral waves emerge at the centre of rotating cores. The spiral is tightly wound with a diffused tail of high amplitude activity that propagates into the rest of the domain and interacts with the other rotating waves. The full spatio-temporal can be seen in Supplemental information 8. Parameter values: *v* = 2, *η*_0_ = 6, *κ*_*v*_ = 0.6, *κ*_*s*_ = 12, *τ* = 1, *α* = 5, *γ* = 0.5.

Increasing *κ*_*v*_ further, results in a ring of incoherence forming between the tightly wound spiral core and the diffused tail (Fig. 10). In addition, radial waves appear to emanate from the outer edge of the ring towards the spiral core. The ring of incoherenc and the radial waves are difficult to distinguish in Fig. 10, but can be seen clearly in Supplemental information 9. Examining the heatmap for the firing rate *R* (top left), we see that the tissue is predominantly silent, with a narrow rotating front of high firing. The temporal dynamics at the green dot, shown in the time course below, reveal highly synchronised burst of activity every 10 to 15 ms. The peaks in firing rate get progressively larger, before reducing again. When an area is silent, the level of synchronisation is still high. This indicates that the neurons are primed to be simultaneously excited when the wave of high activity arrives. If the neurons were desynchronised the wave of high activity would dissipate upon arrival, as the neurons would reach threshold at different times and the peak in firing rate would not be as sharp.

**Figure 10:**
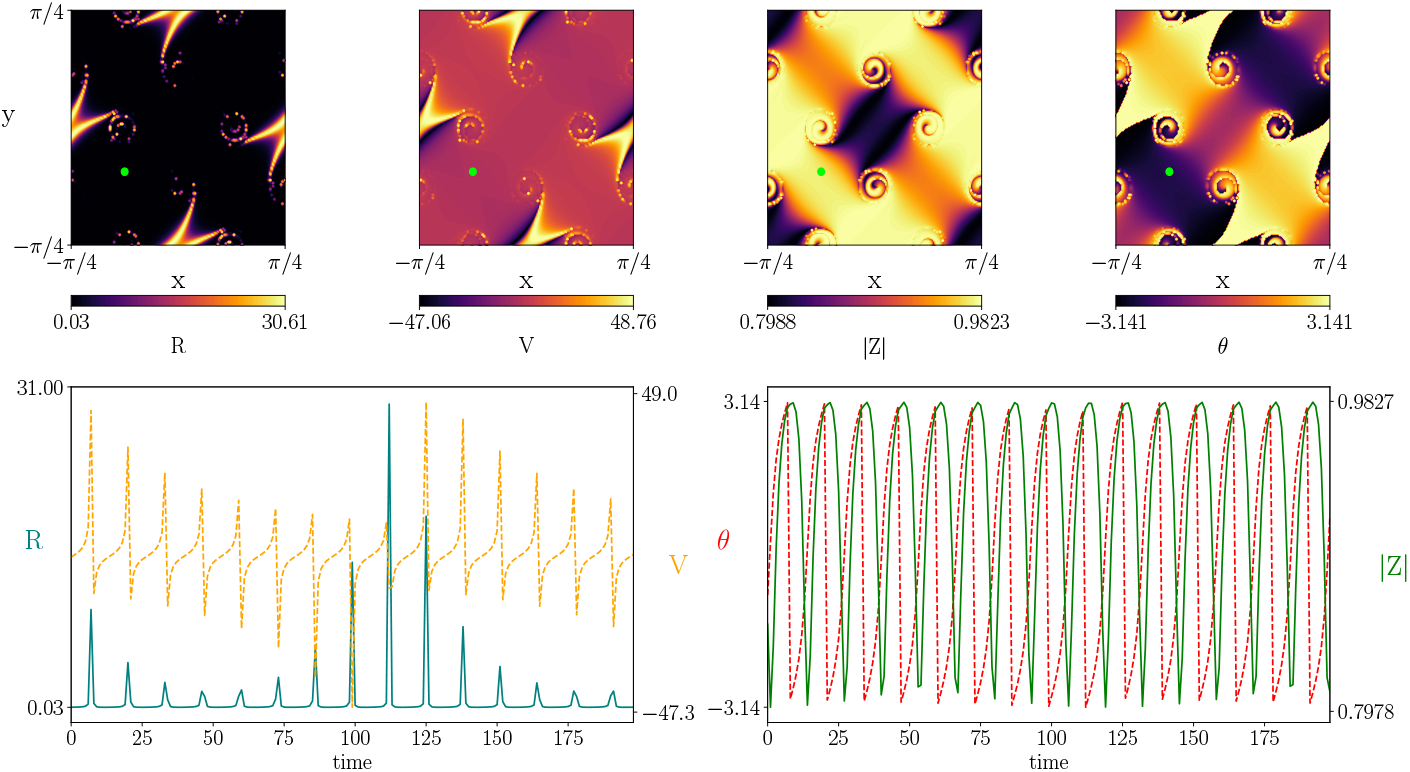
Simulations of the two-dimensional neural field model with large gap junction coupling strength. Note that compared to Fig. 9 the ring of incoherence becomes thicker and radial waves appear to emanate from the outer edge of the ring towards the spiral core. A ring of incoherence exists between the tightly wound spiral and diffused tail, which can be seen more clearly in Supplemental information 9. Parameter values: *v* = 2, *η*_0_ = 6, *κ*_*v*_ = 3.0, *κ*_*s*_ = 12, *τ* = 1, *α* = 5, *γ* = 0.5.

We again note that increasing the gap junction coupling strength increases the level of synchronisation across the tissue. For *κ*_*v*_ = 0.35, the synchrony variable oscillates between 0 and 0.75. For *κ*_*v*_ = 0.6, it oscillates between 0 and 0.88. While for *κ*_*v*_ = 3, the peak synchrony value is almost 1 and the minimum value is roughly 0.80. This supports the hypothesis that gap junction coupling lends to more synchronous activity.

As mentioned in Section 3.1, for a regular wizard hat connectivity kernel (short-range excitation and long-range inhibition) the neural field supports static Turing patterns, periodic bumps of high activity in 1D and a periodic lattice of high-activity spots in 2D. When the Turing bifurcation intersects with the Hopf bifurcation, patterns form at the centre of these localised states. In 2D, patterns of concentric circles can appear within spots when the two bifurcations coincide (Fig. 11). Activity within a localised state can oscillate in time, while the activity in the surround is constant with a low firing rate. These patterns are reminiscent of chimeras [1, 26, 27, 28, 45], as seen in networks of coupled oscillators, where a fraction of the oscillators are phase-locked or silent while the others oscillate incoherently. Note how the peaks in firing rate coincide with peaks in synchrony. However, in the surround synchrony is high, but the firing rate is minuscule. This indicates that the neurons are also synchronised at rest. A video illustrating how these exotic patterns evolve on the entire spatial domain is provided in Supplemental information 10 and the bifurcation diagram is given in Supplemental information 2 panel (c).

**Figure 11:**
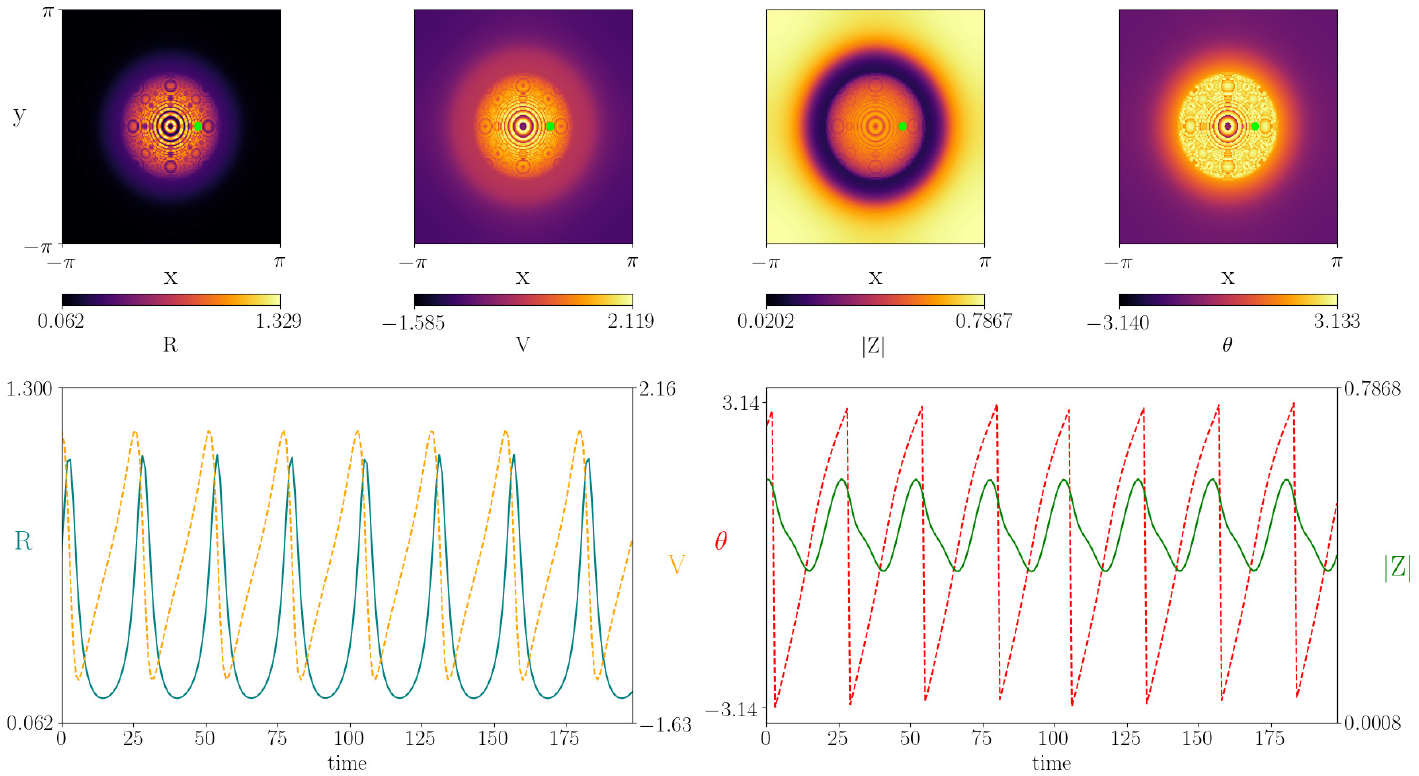
Simulations of the two-dimensional neural field model with shortrange excitation and long-range inhibition, showing the emergence of a spatially localised spot solution (top panel). Note that the core of the spot has a rich temporal dynamics, as indicated in the bottom panel showing the time course for a point within the core (green dot in top panel). A movie showing the full spatio-temporal can be found in Supplemental information 10. Parameter values: *v* = 10.0, *η*_0_ = 0.1, *κ*_*v*_ = 1.0, *κ*_*s*_ = *−*25, *τ* = 1, *α* = 5, *γ* = 0.5.

## 4 Discussion

Mean-field models have proven invaluable in understanding neural dynamics. Although phenomenological in nature, coarse-grained neural mass/field models have proven particularly useful in describing neurophysiological phenomena, such as EEG/MEG rhythms [67], cortical waves [64, 49], binocular rivalry [31, 6], working memory [32] and visual hallucinations [12, 5]. The exclusion of synchrony in standard neural mass/field models prohibits them from describing event-related synchronisation and desynchronisation; the increase and decrease of oscillatory EEG/MEG power due to changes in synchrony within the neural tissue. Here we presented and analysed a recently developed neural mass/field model that incorporates within population synchrony. The main benefit of such a model is that it is derived from a population of interacting spiking neurons. This allows for the inclusion of realistic gap junctions at the cellular level. Gap junctions are known to promote synchrony within neural tissue [61, 4] and the strength of these connections has been linked to the excessive synchronisation driving epileptic seizures [42, 60]. Nonetheless, it is also important to recognise the important effects that the extracellular space has on seizure dynamics, as discussed in [62]. Recent work by Martinet *et al*. [38] has emphasised the usefulness of bringing models to bear on this problem, and coupled the Steyn-Ross neural field model [55] to a simple dynamics for local extracellular potassium concentration. Here, gap-junctions are modelled by appending a diffusive term to a standard neural field and increases in the local extracellular potassium concentration act to decrease the inhibitory-to-inhibitory gap junction diffusion coefficient (to model the closing of gap junctions caused by the slow acidification of the extracellular environment late in seizure). A more refined version of this phenomenological approach would be to replace the Steyn-Ross model with the neural field described here. This would allow a more principled study of how slow changes in the extracellular environment could initiate wave propagation, leading to waves that travel, collide, and annihilate. Indeed, simulations of the next-generation neural field model (without coupling to the extracellular space) have already shown such rich transient dynamics including seizure-like oscillations (and their dependence on the strength of gap-junction coupling). It would be interesting to explore this further, and in particular the transitions whereby spatio-temporal wave patterns are visited in sequence. This has already been the topic of a major modelling study by Roberts *et al*. [49] who considered a variety of more traditional neural mass models in a connectome inspired network using the 998-node Hagmann *et al*. dataset [16] with a single fixed axonal delay. A similar computational study, with a focus on spiral waves and sinks/sources from which activity emanates/converges, could similarly be undertaken using the alternative neural mass model presented here, and with the further inclusion of space-dependent axonal delays. All of the above are topics of ongoing investigation and will be reported upon elsewhere.

## Data availability

Code for running the 1D and 2D neural field simulations can be found at https://github.com/Jamesafross/Neural_Field_with_gaps

## Appendix A Mean field reduction

Consider a heterogeneous network of *N* quadratic integrate-and-fire neurons with voltage *v*_*i*_ and both gap-junction and synaptic coupling:

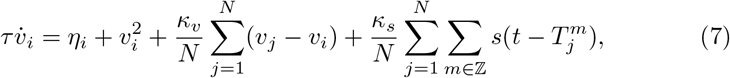

*i* = 1, …, *N*, *v*_*r*_ ≤ *v*_*i*_ ≤ *v*_th_ Here, the *m*th firing time of the *j*th neuron is defined implicitly by 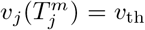. The network nodes are subject to reset: *v*_*i*_ → *v*_r_ at times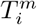. The strengths of gap-junction and synaptic coupling are *κ*_*v*_ and *κ*_*s*_ respectively. The function *s*(*t*) represents the shape of a post synaptic response (to a delta-Dirac spike) and will be taken to be the Green’s function of a linear differential operator *Q*. For an alpha-function *s*(*t*) = *α*^2^*t* exp(*αt*)*H*(*t*), where *H* is a Heaviside function, *Q* = (1 + *α^−^*^1^d*/*d*t*)^2^, whilst for an exponential response *s*(*t*) = *α* exp(−*αt*)*H*(*t*), *Q* = (1 + *α*^−1^*d*/*dt*). In (7) the *η*_*i*_ are random variables drawn from a Lorentzian distribution:

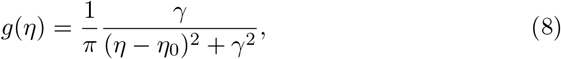

with mean *η*_0_ and half-width *γ*. For simplicity we shall take *v*_*r*_ → −*∞* and *v*_*th*_ → *∞*.

To derive the mean field equations we follow closely the exposition by Mont-brió *et al*. [39]. Consider the thermodynamic limit *N* → *∞* with a distribution of voltage values *ρ*(*v*|*η*, *t*). The continuity equation for *ρ* is

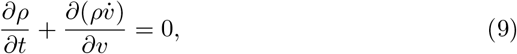

where

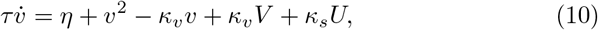

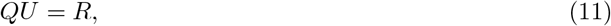

and

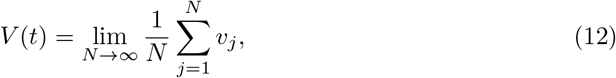

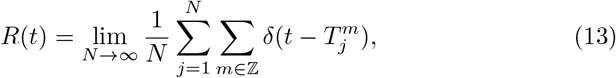

which represent the average voltage and population firing rate respectively. We now assume a solution *ρ*(*v η, t*) of the form

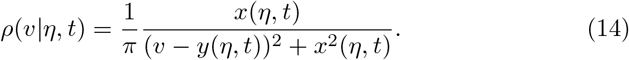

For a fixed *η* the firing rate *r*(*η, t*) can be calculated as 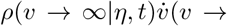 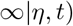, from which we may establish that

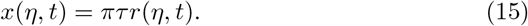

By exploiting the structure of (14), with poles at *v*_*±*_ = *y* ± *ix*, a contour integration shows that

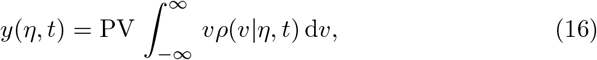

where PV denotes the Cauchy principal value. After averaging over the distribution of single neuron drives given by (8) we obtain

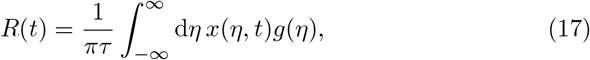

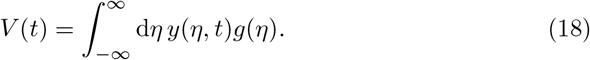

For fixed *η*, substitution of (14) into the continuity equation and balancing powers of *v* shows that *x* and *y* obey two coupled differential equations that can be written as

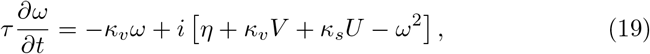

where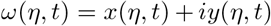. After evaluating the integrals in (42) and (18) using contour integration (and using the fact that *ρ* has poles at *η*_*±*_ = *η*_0_ ± *iγ*) the coupled equations for (*R, V*) can be found as

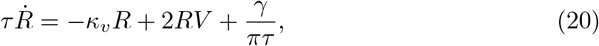

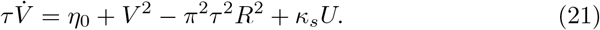

The complex quantity *W* = *πτ R* + *iV* is known to be related to the Kuramoto order parameter *Z* by the conformal map [39]:

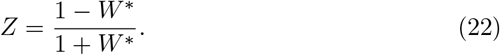

The evolution equation for *Z* is given by the complex differential equation

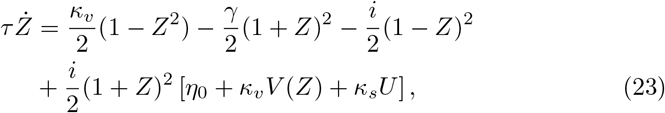

where *QU* = *R*(*Z*) and

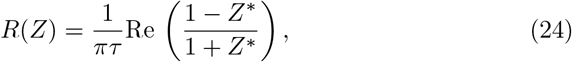

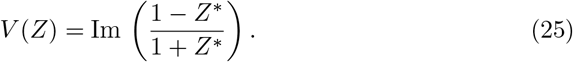

## Appendix B Interacting sub-populations

Consider an excitatory population labelled by *E* coupled to an inhibitory one labelled by *I*. In this case there are four distinct synaptic inputs with connection strengths 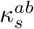, *a, b ∈ {E, I}*, with 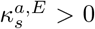 and 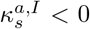. Each population has a background drive drawn from a Lorentzian with mean 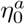 and half-width *γ*_*a*_, *a* ∈ {*E, I*}. Generalising the mean field model derived in section Appendix A, we have that

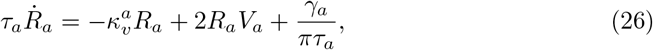

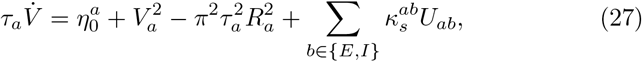

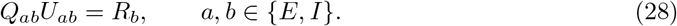

For a second order synapse with time-scale 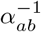 we would set

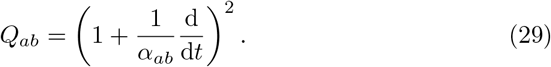

## Appendix C Brain wave equation

A simple continuum model for an effective single population dynamics can be written in the form

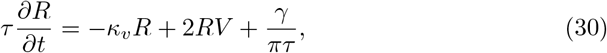

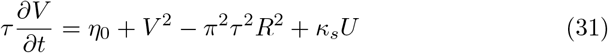

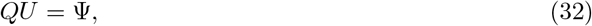

where ψ = *w* ꕕ *R*. The symbol ꕕ is used to describe spatial interaction within the neural field model, while *w* represents structural connectivity. For example, in the plane we might consider (*R, V, U*) = (*R*(**r***, t*)*, V* (**r***, t*)*, U* (**r***, t*)), with **r***∈* ℝ^2^ and *t ≥* 0 with

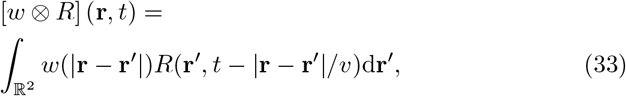

where *v* represents the speed of an action potential. We note that (33) can be written as a convolution:

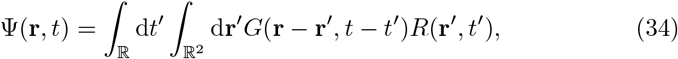

where *G*(*r, t*) = *w*(*r*)*δ*(*t* − *r/v*). For certain choices of *w* it is possible to exploit this convolution structure to obtain a PDE model, often referred to as a *brain-wave equation* [43, 24].

For the choice of an inverted balanced wizard hat function with *w*(*r*) = (*r/*2 − 1)*e*^−*r*/^(2*π*) this approach yields the following brain-wave equation:

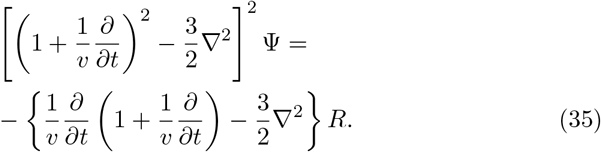

Note that (35) is only strictly valid for describing long-wavelength solutions. In one spatial dimension and using *w*(*x*) = (|*x*| − 1)*e*^−|*x*|^ the brain-wave PDE is

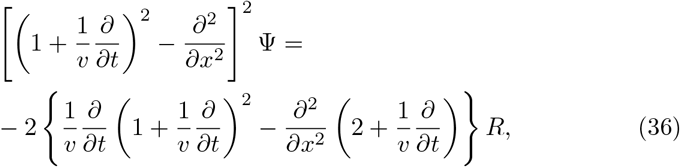

and is an exact reduction of ψ = *w* ꕕ *R* [59].

## Appendix D Turing instability analysis

Consider the homogeneous steady state of (35) given by (*U* (**r***, t*), ψ(**r***, t*)*, R*(**r***, t*)*, V* (**r***, t*)) = (0, 0, *R*_0_*, V*_0_) where (*R*_0_*, V*_0_) are given by the simultaneous solution of the algebraic equations

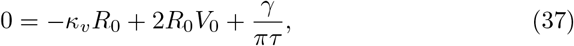

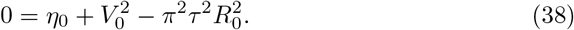

We linearise around the steady state and consider perturbations of the form (*U* (**r***, t*), ψ(**r***, t*)*, R*(**r***, t*), 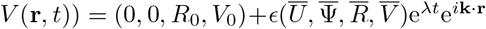 for *ϵ*: ≪ 1 and 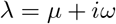. Substitution into (35) and working to first order in *ϵ* gives the linear relationship

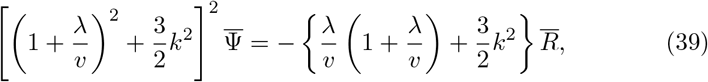

where *k* = *|***k***|*. A linearisation for the dynamics of (*R, V*) gives

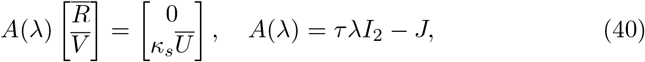

where *I*_2_ is the 2 *×* 2 identity matrix and *J* is the Jacobian

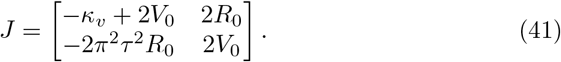

We may solve (40) using Cramer’s rule to yield

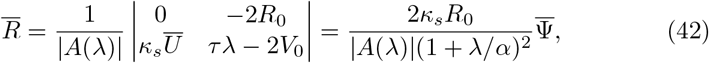

where we have used the fact that 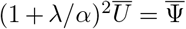 (from (32)). Substitution of 42 into (39) and demanding a non-trivial solution for 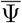 leads to the condition Ɛ(*λ, k*) = 0, where

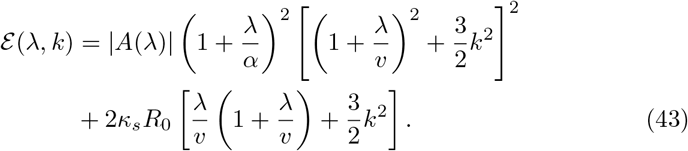

Thus, the continuous spectrum *λ* = *λ*(*k*) is given by the roots of an eight order polynomial.

A similar analysis of (36) gives

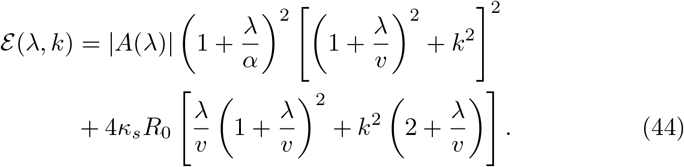

The system undergoes a bifurcation when a branch of solutions *λ*(*k*) to Ɛ(*λ*, *k*) = 0 touches the imaginary axis, *µ*(*k*_*c*_) = 0, where *k*_*c*_ is the critical wave number. By the implicit function theorem, this occurs when

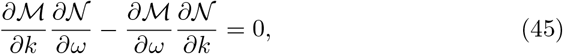

where 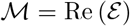 and 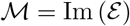.

A Hopf bifurcation of the spatially uniform can be found by solving 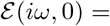 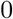 for 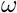. A static Turing bifurcation is found by solving *Ɛ* (0, *k*_*c*_) = 0 and (45) for *k*_*c*_ non-zero. While a dynamic Turing-Hopf bifurcation is found by solving 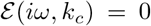 and (45) for non-zero 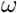 and *k*_*c*_. Interesting patterns tend to emerge when a Hopf and Turing-Hopf intersect at a codimension-2 point. Such a bifurcation can be found by solving 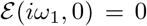, 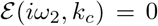 and (45) simultaneously for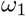, 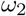 and *k*_*c*_.

## List of Supplemental information

### Supplemental information 1

Chimeras in the one-dimensional neural field model. (a) Bifurcation diagram for a standard wizard hat coupling kernel. (b) Simulation close to the Turing and Hopf curves *η*_0_ = − 0.1 and *κ*_*v*_ = 1. Other parameter values: *v* = 2, *κ*_*s*_ = − 15, *τ* = 1, *α* = 3, *γ* = 0.5.

### Supplemental information 2

Turing analysis for two-dimensional neural field model. (a) Hopf and Turing-Hopf curves for an inverted wizard hat coupling kernel (*κ*_*s*_ = 12) with *η*_0_ = 0.5. (b) Hopf and Turing-Hopf curves when *η*_0_ is increased to 0.6 and *κ*_*s*_ remains unchanged. (c) Bifurcation diagram for a standard wizard hat coupling kernel (*κ*_*s*_ = *−*25), with *v* = 10. Other parameter values: *τ* = 1, *α* = 5, *γ* = 0.5.

### Supplemental information 3

Bulk oscillations appear in the 2D neural field model when we simulate close to the Hopf bifurcation. The entire tissue oscillates coherently and there are no spatial patterns. Here we show the oscillations for the synchrony variable *|Z|*. Parameter values: *κ*_*v*_ = 0.1, *v* = 4, *η*_0_ = 6, *κ*_*s*_ = 12, *τ* = 1, *α* = 5, *γ* = 0.5.

### Supplemental information 4

When close to the Turing-Hopf bifurcation, perturbing the 2D neural field model with horizontal bars of high activity results in planar waves. A high activity source forms at the centre of the domain and the waves propagate up and down from it. The evolution of the synchrony variable *|Z|* is shown here. Parameter values: *κ*_*v*_ = 0.35, *v* = 2, *η*_0_ = 6, *κ*_*s*_ = 12, *τ* = 1, *α* = 5, *γ* = 0.5.

### Supplemental information 5

The uniform steady state of the 2D neural field model was perturbed with a Gaussian to initiate radial waves. The temporal evolution of the waves is shown for the synchrony variable |*Z*|. A source of high activity emerges at the centre of the domain and the waves emanate from it. Periodic boundary conditions were used, and as such, the waves interfere when they reach the edge of the domain. Parameter values: *κ*_*v*_ = 0.35, *v* = 2, *η*_0_ = 6, *κ*_*s*_ = *−*15, *τ* = 1, *α* = 3, *γ* = 0.5.

### Supplemental information 6

Standing waves emerge in the 2D neural field model when we are close to the Turing-Hopf bifurcation and the uniform steady state is perturbed with a spatially periodic lattice pattern. We show the temporal dynamics of the synchrony variable *|Z|* in this movie. The minima and maxima alternate in time, but the lattice pattern remains unchanged. Parameter values: *κ*_*v*_ = 0.35, *v* = 2, *η*_0_ = 6, *κ*_*s*_ = 12, *τ* = 1, *α* = 5, *γ* = 0.5.

### Supplemental information 7

Simulation of the 2D neural field model corresponding to the weak gap junction coupling regime shown in Fig. 8. The temporal evolution of synchrony variable |*Z*| is shown here. The steady state was perturbed with a spatially periodic lattice and rotating waves emerge. We observe source and sink dynamics were the waves interact with one another. Parameter values: *κ*_*v*_ = 0.35, *v* = 2, *η*_0_ = 6, *κ*_*s*_ = 12, *τ* = 1, *α* = 5, *γ* = 0.5.

### Supplemental information 8

Spatio-temporal dynamics of the synchrony variable |*Z*| in the 2D neural field model with an intermediate value of gap junction coupling strength. The simulation corresponds to Fig. 9, where tightly wound spirals appear the centre of the rotating waves. Parameter values: *v* = 2, *η*_0_ = 6, *κ*_*v*_ = 0.6, *κ*_*s*_ = 12, *τ* = 1, *α* = 5, *γ* = 0.5.

### Supplemental information 9

Simulation the 2D neural field model with strong gap junction coupling, as seen in Fig. 10. We show the temporal dynamics of the synchrony variable |*Z*|, which varies between ~ 0.8 and 1. A ring of incoherence emerges between the tightly wound spiral and diffused tail, with radial waves propagating from the edge of the ring toward the centre of the rotating core. Parameter values: *v* = 2, *η*_0_ = 6, *κ*_*v*_ = 3.0, *κ*_*s*_ = 12, *τ* = 1, *α* = 5, *γ* = 0.5.

### Supplemental information 10

Simulation of the chimera dynamics in the 2D neural field model with a regular wizard hat coupling kernel. Movie corresponds to Fig. 11 and shows the dynamics of the synchrony variable *|Z|*. Parameter values: *v* = 10.0, *η*_0_ = 0.1, *κ*_*v*_ = 1.0, *κ*_*s*_ = *−*25, *τ* = 1, *α* = 5, *γ* = 0.5.

## Notes

https://github.com/Jamesafross/Neural_Field_with_gaps

### Competing Interest Statement

The authors have declared no competing interest.

