## Supplementary figures and images for "Mean-field models for EEG/MEG: from oscillations to waves"

### Supplemental information 1

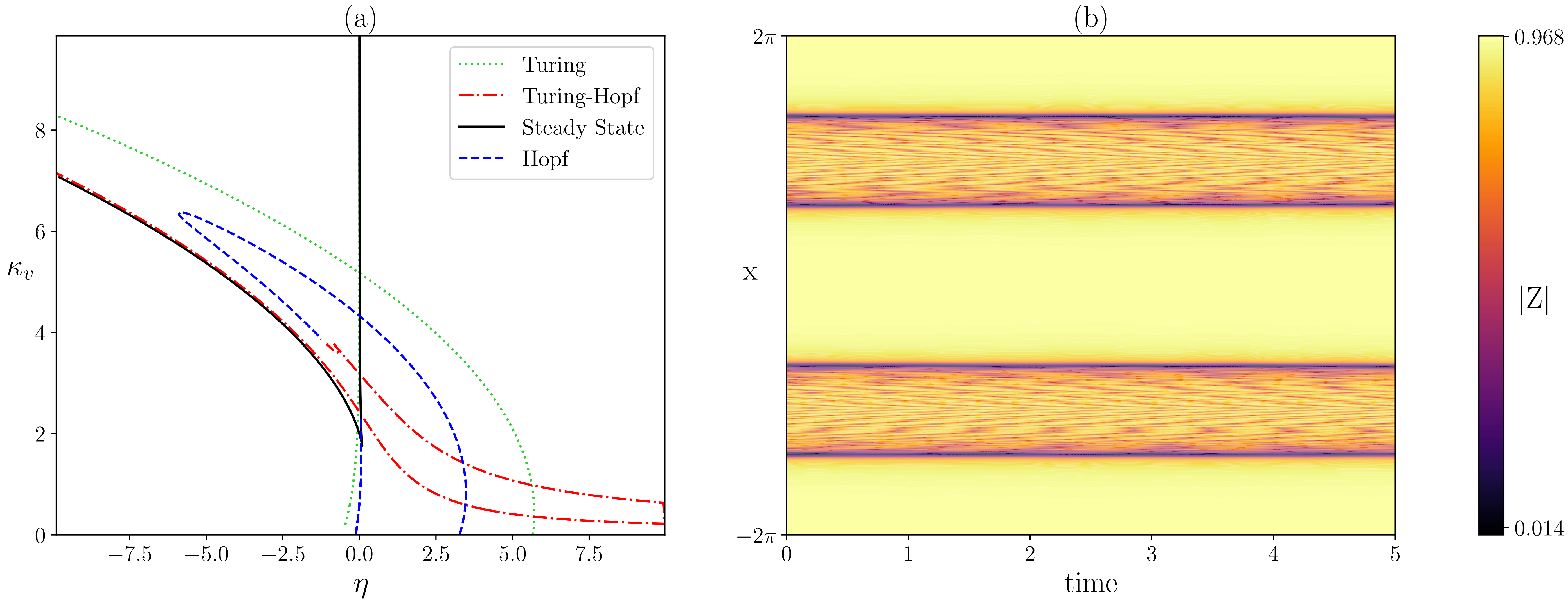

### Supplemental information 2

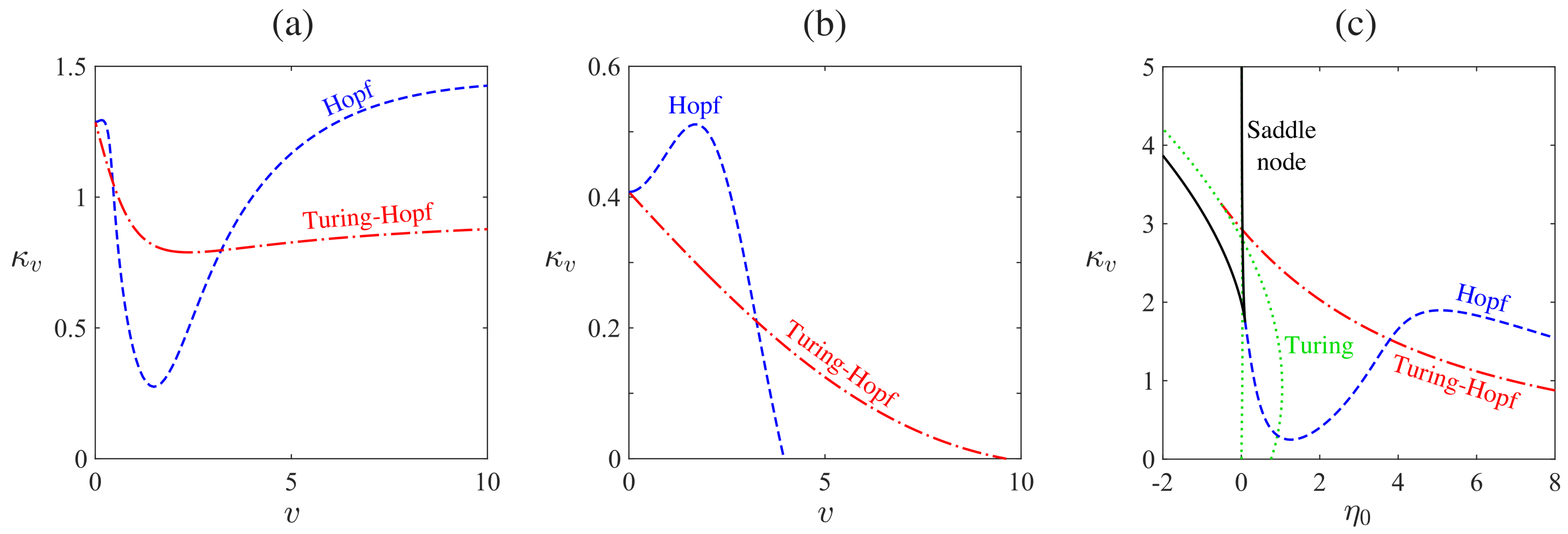
